# The role of Nef in the long-term persistence of the replication-competent HIV reservoir in South African women

**DOI:** 10.1101/2024.11.01.621615

**Authors:** Sherazaan D. Ismail, Shorok Sebaa, Bianca Abrahams, Martha C. Nason, Mitchell J. Mumby, Jimmy D. Dikeakos, Sarah B. Joseph, Matthew Moeser, Ronald Swanstrom, Nigel Garrett, Carolyn Williamson, Thomas C. Quinn, Melissa-Rose Abrahams, Andrew D. Redd

## Abstract

HIV-1 Nef mediates immune evasion and viral pathogenesis in part through downregulation of cell surface cluster of differentiation 4 (CD4) and major histocompatibility complex class I (MHC-I) on infected cells. While Nef function of circulating viral populations found early in infection has been associated with reservoir size in early-treated cohorts, there is limited research on how its activity impacts reservoir size in people initiating treatment during chronic infection. In addition, there is little research on its role in persistence of viral variants during long-term antiretroviral therapy (ART). Phylogenetically distinct *nef* genes (n=82) with varying estimated times of reservoir entry were selected from viral outgrowth variants stimulated from the reservoir of South African women living with HIV who initiated ART during chronic infection (n=16). These *nef* genes were synthesized and used in a pseudovirus infection assay that measures CD4 and MHC-I downregulation via flow cytometry. Downregulation measures were compared to the size of the replication-competent viral reservoir (RC-VR), estimated by quantitative viral outgrowth assay (QVOA) at 5 years after treatment initiation, as well as proviral survival time. Maximum Nef-mediated MHC-I downregulation was significantly associated with RC-VR size (p=0.034), but this association was not observed for CD4 downregulation. Conversely, we did not find a consistent association between intraparticipant MHC-I or CD4 downregulation and the variant timing of entry into the reservoir. These data support a role for Nef-mediated MHC-I downregulation in determining RC-VR size, but more work is needed to determine Nef’s role in the survival of individual viral variants over time.

**Author summary:** Understanding the HIV reservoir, including viral determinants of reservoir formation and maintenance, is key for the rational design of cure interventions. In addition, for cure interventions to be equitable, it needs to be broadly applicable. While African women bear the greatest burden of HIV globally, most cure research has been performed in men living in the global North. Our study aims to elucidate attributes of the virus that contribute to reservoir dynamics in South African women on ART. We investigated the ability of the HIV accessory protein Nef to reduce the cell surface levels of two cellular proteins, CD4 and MHC-I, and compared this downregulation capacity with the size of the HIV reservoir and survival of cells infected with a given viral isolate. We found a positive association between an individual’s measured reservoir size and MHC-I downregulation, but not CD4 downregulation. There was little evidence for a survival benefit for stronger Nef MHC-I reduction, but more research is needed on this subject. These data support earlier work and suggest that Nef’s interaction with MHC-I may be a target to restrict the latent reservoir and inform alternate cure strategies.

## Introduction

Human Immunodeficiency Virus (HIV) is effectively managed with antiretroviral therapy (ART), which suppresses viral replication to below detectable levels. However, viral eradication in people living with HIV (PLWH) is impeded by the early formation of a stable reservoir, primarily in resting CD4^+^ T cells [1–3]. While the majority of proviruses in this viral reservoir are defective and cannot produce infectious virus [4], the remaining small percentage of intact proviruses can reactivate upon ART interruption, resulting in viral recrudescence [5–9]. Major efforts have been undertaken to understand the formation of the viral reservoir and factors affecting the persistence of latent proviruses. While the pool of infected reservoir cells is established very early [10], and early ART initiation has been shown to restrict reservoir size [11–13], some studies have found that ≥60% of viral variants persisting during long-term ART are those present immediately preceding ART initiation [14–18], regardless of the duration of untreated infection. However, there is a paucity of information on the underlying host and viral mechanisms of reservoir establishment and maintenance over time.

One viral factor that has been associated with HIV reservoir dynamics is the accessory protein, Nef. Nef is a polyfunctional protein that mediates immune evasion, infectivity, and pathogenicity through disruption of host cell antiviral activity [19–21]. Two key functions of Nef include downregulation of cluster of differentiation (CD4) and major histocompatibility complex class I (MHC-I) from the surface of infected cells. CD4 downregulation restricts natural killer (NK) cell clearance of infected cells through antibody-dependent cytotoxicity (ADCC) [22], while MHC-I downregulation prevents infected cell recognition by cytotoxic T lymphocytes (CTLs) [23,24]. There is evidence that intact Nef can be expressed during long-term ART [25–27], even if the remainder of the provirus is defective [28–30] and that these intact Nef proteins can mediate MHC-I downregulation and preclude cells from CTL clearance [28,29]. In addition, the ability of Nef to downregulate MHC-I *in vitro* has been associated with *in vivo* reservoir size in men on ART for ∼one year, treated during early infection [31]. This relationship was not present between reservoir size and CD4 downregulation by Nef. Therefore, the capacity of Nef to downregulate MHC-I may play a role in sustained immune evasion on long-term ART, potentially aiding viral persistence.

We hypothesized that the degree of MHC-I, but not CD4, downregulation by Nef would be associated with a greater frequency of latently infected cells, as well as the survival of variants in the replication-competent HIV reservoir (RC-VR; here, defined as infectious units per million T cells as measured by QVOA).

## Results

*Nef* genes sequenced from outgrowth viruses (OGVs) from South African women living with HIV (n=16) were selected for functional analyses [16,18] (**Table 1**). All *nef* sequences were predicted to be subtype C using the Geno2Pheno subtyping tool (**Supplementary Table S1**) [32].

**Table 1.**
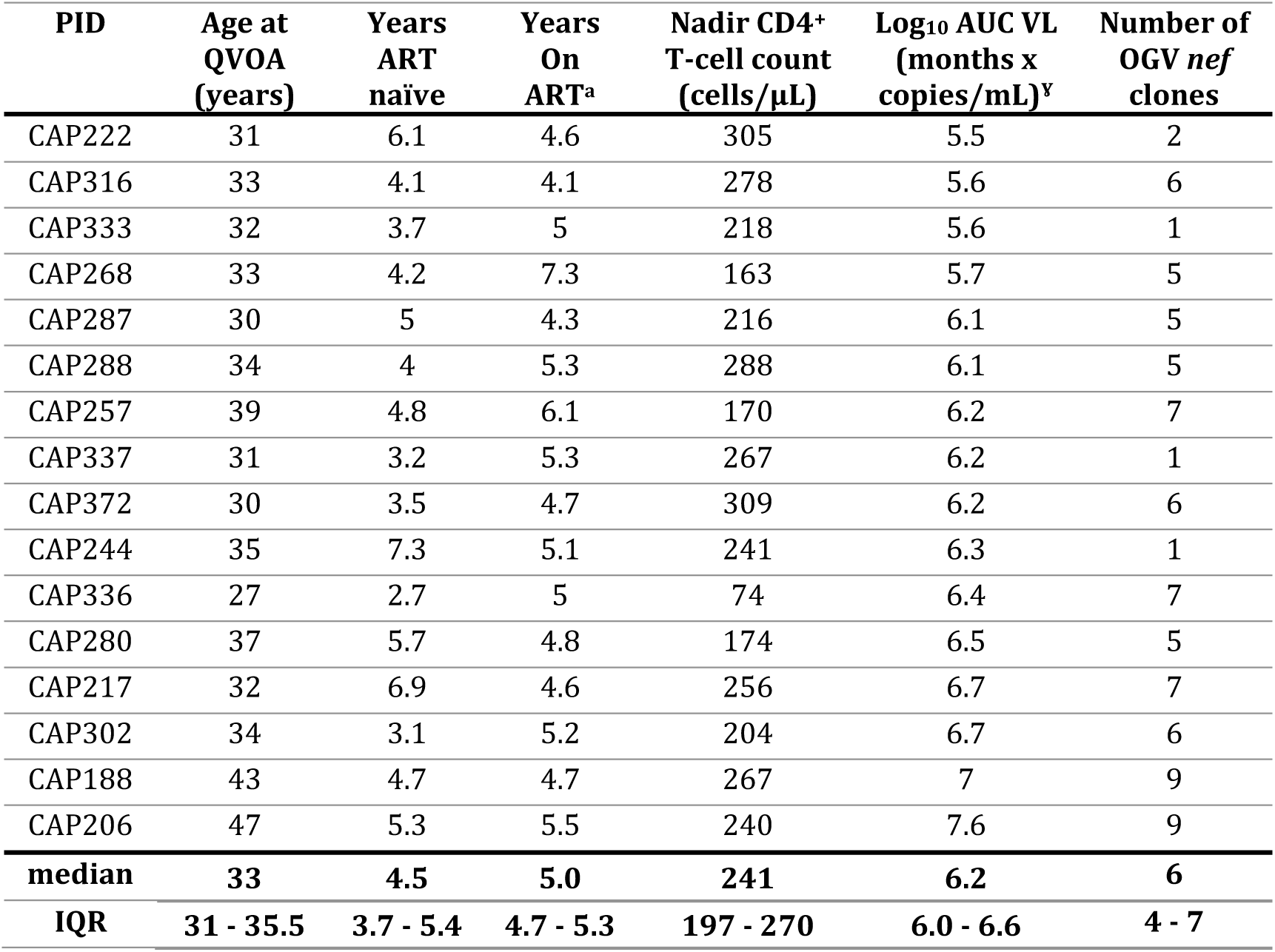
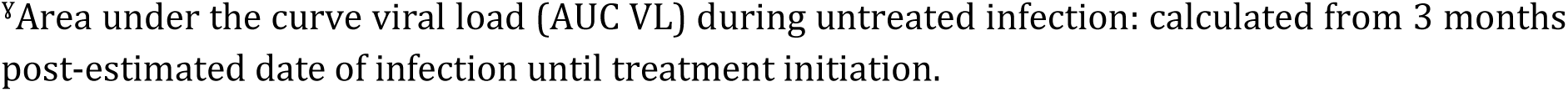
Participant information.

| PID | Age at QVOA (years) | Years ART naïve | Years On ART<sup>a</sup> | Nadir CD4<sup>+</sup> T-cell count (cells/μL) | Log<sub>10</sub> AUC VL (months x copies/mL)<sup>y</sup> | Number of OGV *nef* clones |
| --- | --- | --- | --- | --- | --- | --- |
| CAP222 | 31 | 6.1 | 4.6 | 305 | 5.5 | 2 |
| CAP316 | 33 | 4.1 | 4.1 | 278 | 5.6 | 6 |
| CAP333 | 32 | 3.7 | 5 | 218 | 5.6 | 1 |
| CAP268 | 33 | 4.2 | 7.3 | 163 | 5.7 | 5 |
| CAP287 | 30 | 5 | 4.3 | 216 | 6.1 | 5 |
| CAP288 | 34 | 4 | 5.3 | 288 | 6.1 | 5 |
| CAP257 | 39 | 4.8 | 6.1 | 170 | 6.2 | 7 |
| CAP337 | 31 | 3.2 | 5.3 | 267 | 6.2 | 1 |
| CAP372 | 30 | 3.5 | 4.7 | 309 | 6.2 | 6 |
| CAP244 | 35 | 7.3 | 5.1 | 241 | 6.3 | 1 |
| CAP336 | 27 | 2.7 | 5 | 74 | 6.4 | 7 |
| CAP280 | 37 | 5.7 | 4.8 | 174 | 6.5 | 5 |
| CAP217 | 32 | 6.9 | 4.6 | 256 | 6.7 | 7 |
| CAP302 | 34 | 3.1 | 5.2 | 204 | 6.7 | 6 |
| CAP188 | 43 | 4.7 | 4.7 | 267 | 7 | 9 |
| CAP206 | 47 | 5.3 | 5.5 | 240 | 7.6 | 9 |
| **median** | **33** | **4.5** | **5.0** | **241** | **6.2** | **6** |
| **IQR** | **31 - 35.5** | **3.7 - 5.4** | **4.7 - 5.3** | **197 - 270** | **6.0 - 6.6** | **4 - 7** |

<sup>y</sup>Area under the curve viral load (AUC VL) during untreated infection: calculated from 3 months post-estimated date of infection until treatment initiation.

The women in this group were living with HIV for a median of 4.5 years prior to ART initiation and were on ART for a median of 5 years prior to sampling OGVs. As part of this earlier analysis, the estimated time of entry for each given OGV into the reservoir was determined using phylogenetic approaches [16,18]. A total of 82 *nef* sequences (median = 6 per participant) were selected based on phylogenetic clustering on an amino acid Maximum Likelihood tree and entry timing estimates (**Figure S1**). The corresponding *nef* genes were synthesized and cloned into a single-round infection based pseudovirus (PSV) reporter system for examining *in vitro* Nef-mediated CD4 and MHC-I downregulation as previously described [33]. PSVs represented viruses estimated to enter the reservoir over a range of 30 weeks to 6.3 years post-infection (**Figure 1**).

**Figure 1.**
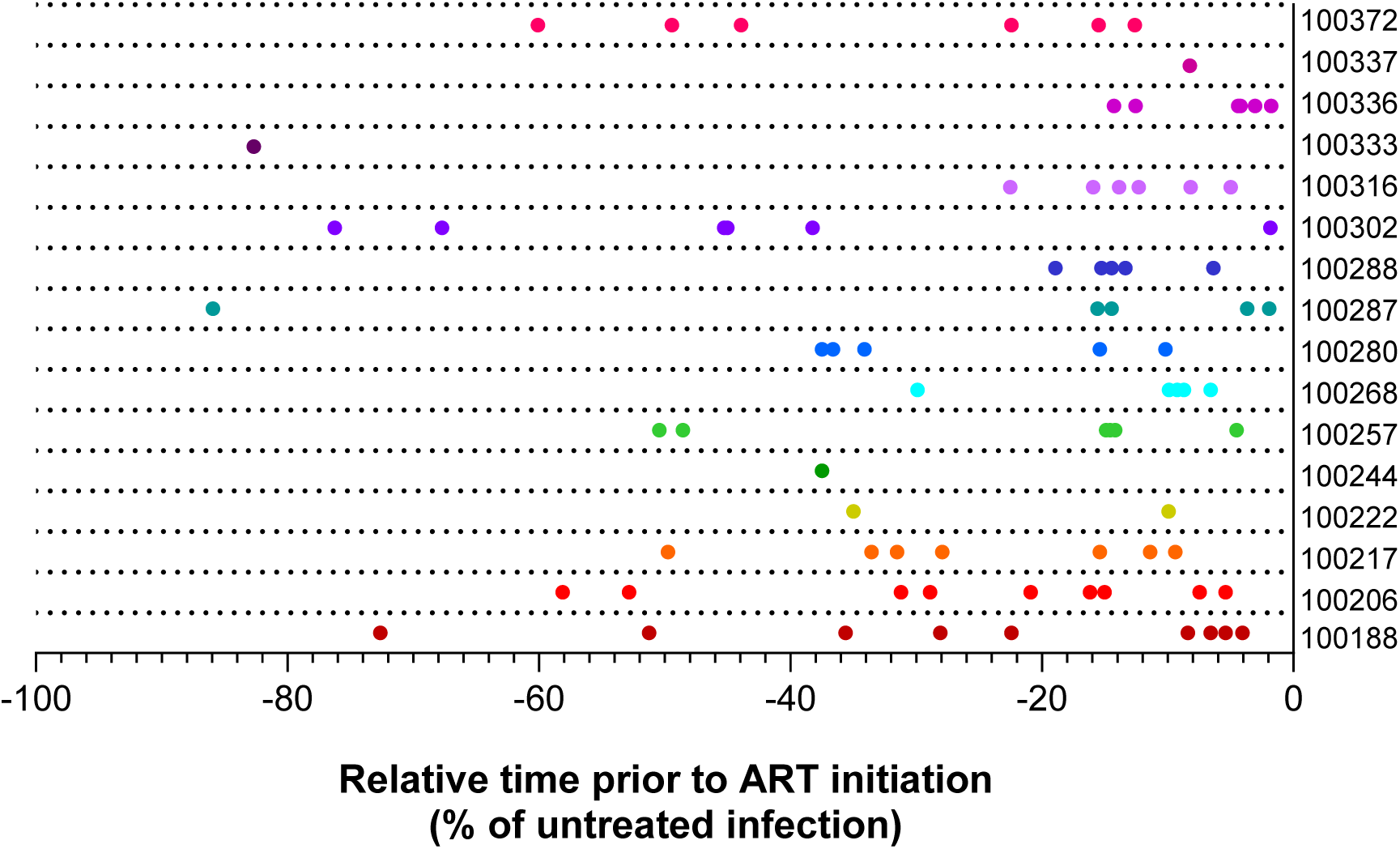
Estimated timing of reservoir entry distribution of Nefs selected for functional analyses. The timeline shows the estimated timing of entry (x-axis) into the reservoir of each selected outgrowth virus Nef relative to the estimated time of infection (-100%) and the relative start of ART (0%) for each participant. Data points are coloured by participant.

The diversity of *nef* sequences within each participant was evaluated by comparing pairwise distances in MEGA 11 [34]. Mean pairwise DNA distances ranged from 0.0079 to 0.0266, while maximum pairwise distances ranged from 0.0079 to 0.0549 **(Figure 2, and S1**).

**Figure 2.**
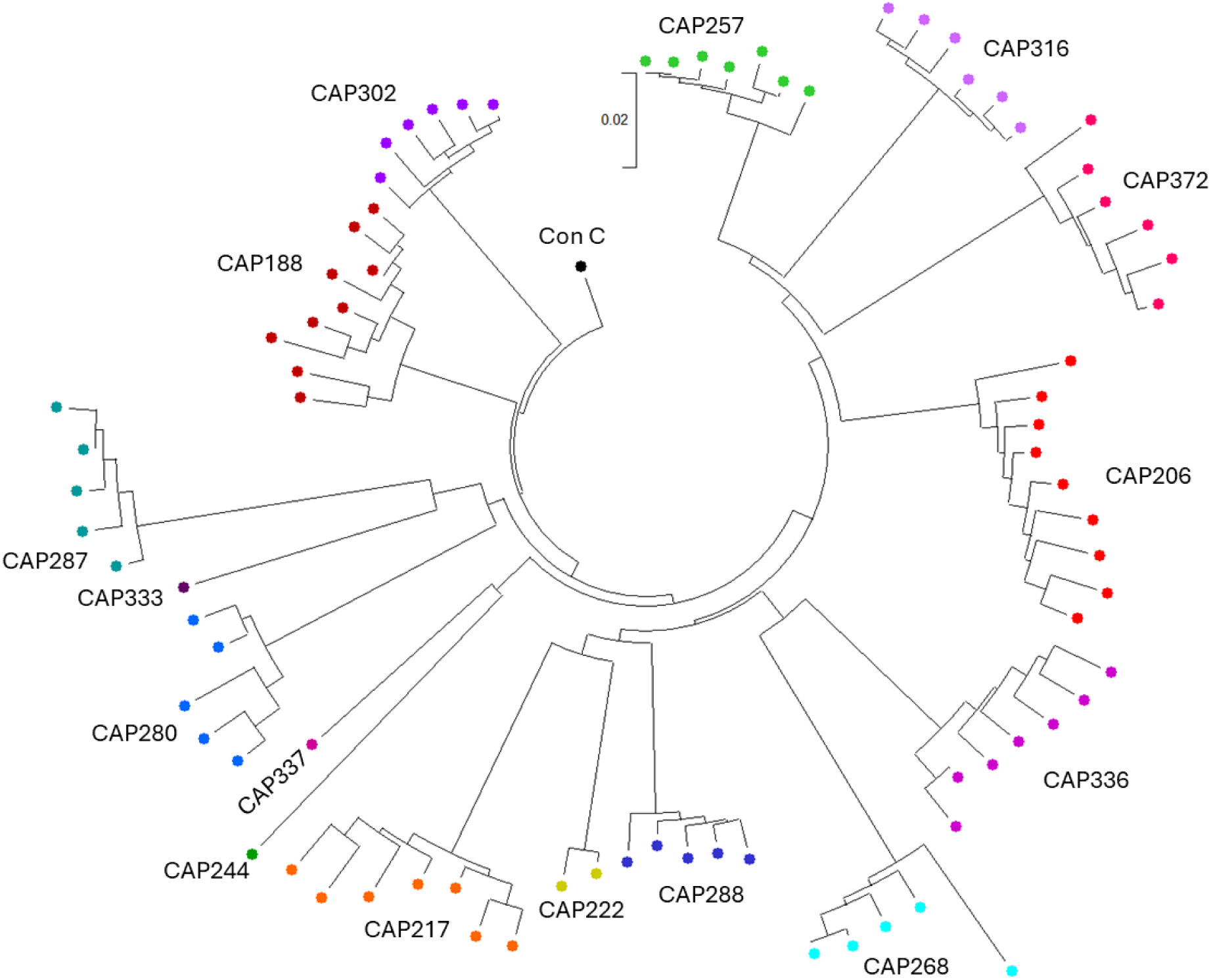
The phylogenetic relationship between *nef* gene sequences examined in this study. OGV *nef* nucleotide sequences were codon-aligned (Clustal W in MEGA11) and the tree was constructed using the Neighbour-Joining method [35] and rooted to the Consensus C *nef* nucleotide sequence obtained from the LANL database (https://www.hiv.lanl.gov/). The optimal tree is shown. This analysis involved 83 nucleotide sequences including Consensus C *nef*. Codon positions included were 1st+2nd+3rd+Noncoding. All ambiguous positions were removed for each sequence pair (pairwise deletion option). There was a total of 670 positions in the final dataset. Evolutionary analyses were conducted in MEGA11 [34].

Nef function was assessed as the downregulation of cell surface CD4 and MHC-I after infection of SUPT1 cells for 48h with participant-specific Nef PSVs. Five to 20 replicate infections were performed per PSV (median = 9). MHC-I downregulation ranged from 2.53 to 5.19 times the ΔNef control (mean=3.92) and from 0.60 to 11.69 times the ΔNef control for CD4 downregulation (mean=2.74) (**Figure 3**). A within-participant comparison for those individuals with more than one *nef* clone (n=13) indicated that only four individuals had significant differences in CD4 downregulation function within the selected OGV Nef proteins, while nine individuals had significant differences in MHC-I downregulation function (Kruskal-Wallis H test P<0.05; indicated by black lines on the right of each plot in **Figure 3**).

**Figure 3.**
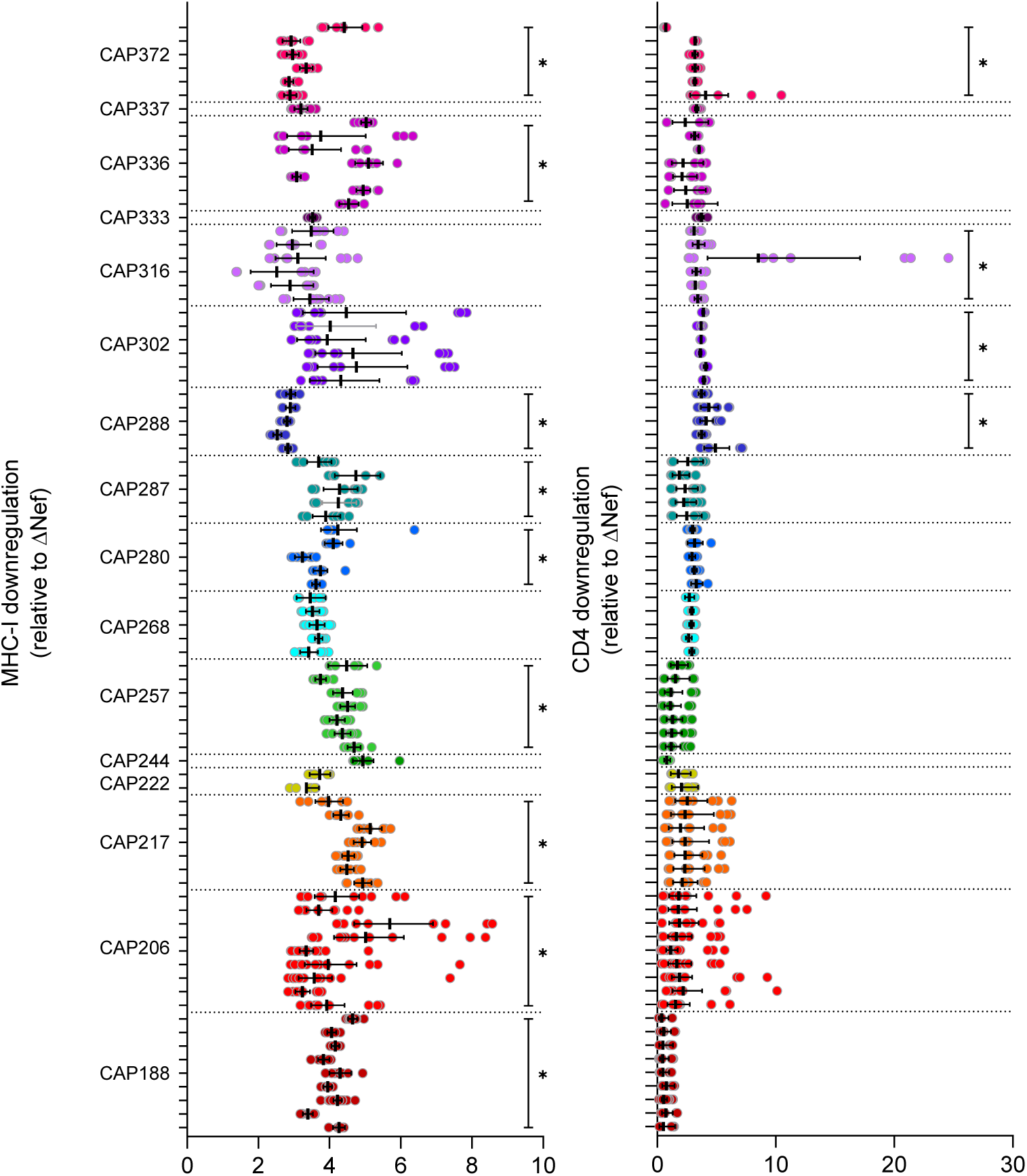
Nef-mediated MHC-I and CD4 downregulation. Individual replicates for Nef-mediated downregulation of MHC-I (left) or CD4 (right) compared to a Nef-deleted control PSV (ΔNef). Bars represent the geometric mean downregulation after infection of SUPT1 cells with different Nef-typed PSVs. Error bars represent the 95% CI. Each participant is represented by a different colour (in descending PID order from top to bottom), with clones from the same participant grouped together on the figure. Black lines on the right of each plot indicate individuals where within-participant Nef function differed significantly after Kruskal-Wallis H Tests (non-parametric one-way ANOVA) were performed with a P-value cut-off of < 0.05 considered significant.

We initially evaluated whether there was a relationship between Nef-mediated CD4 or MHC-I downregulation and RC-VR size. For these cross-sectional analyses, we evaluated within-participant maximal CD4 or MHC-I downregulation (i.e. the clone with maximal downregulation activity). There was a significant positive relationship between reservoir size and maximal MHC-I downregulation (p=0.0344; slope=0.204; **Figure 4A**), but not maximal CD4 downregulation (p=0.6302; **Figure 4B**). However, when adjusted for age, nadir CD4 count, or AUC VL, the relationship between maximal MHC-I downregulation and reservoir size was no longer significant.

**Figure 4.**
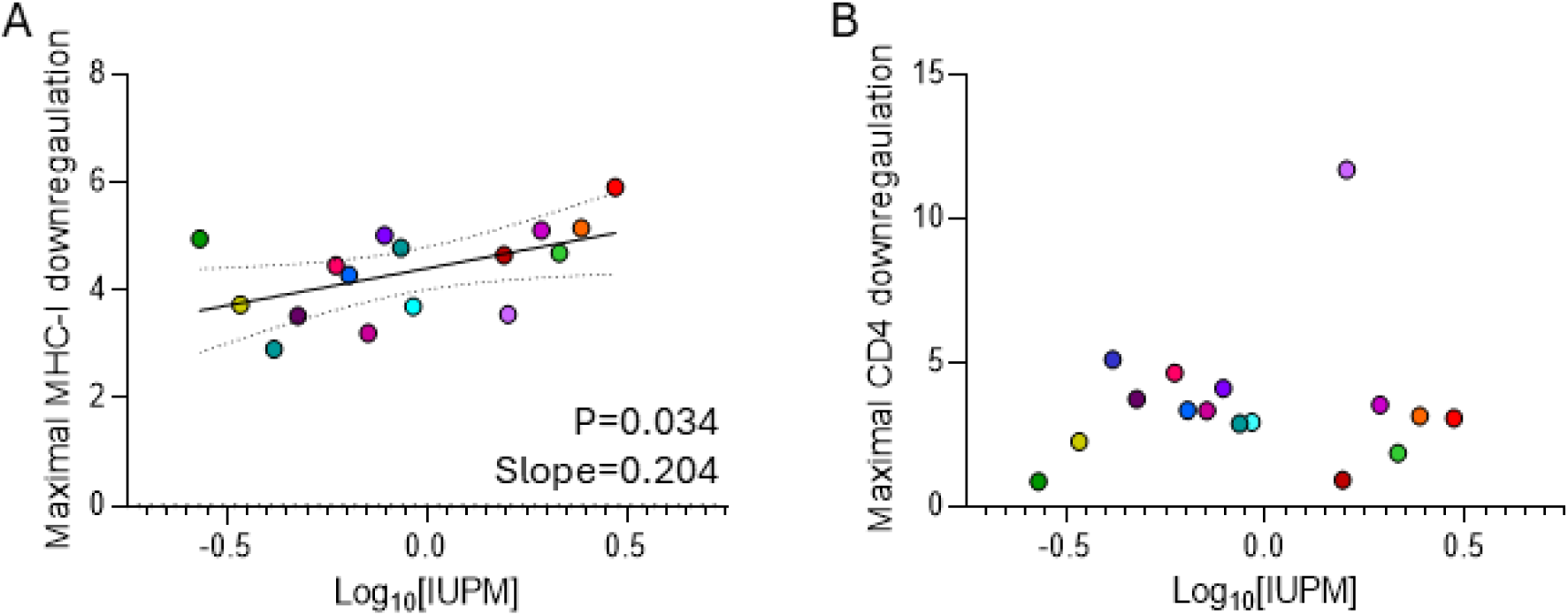
The relationship between the frequency of latently infected cells containing replication-competent HIV (reservoir size) and maximal MHC-I (A) or CD4 (B) downregulation by participant-specific Nef. Reservoir size is represented as the number of infectious units per million CD4^+^ T cells (IUPM). Linear regression best fit lines and 95% CI were plotted for significant linear relationships.

We next wanted to determine whether Nef function is a correlate of viral persistence for individual proviruses in the latent reservoir, hypothesizing that stronger downregulation of MHC-I by Nef facilitates longer proviral survival. Using the estimated timing of entry of viral variants in the latent reservoirs of these women [16,18], we calculated the proviral survival time for each variant (combined time from estimated reservoir entry to QVOA sampling time while on ART). Mixed effects regression (without any co-variates) showed no relationship between proviral survival time and either geometric mean CD4 (p=0.221) or geometric mean MHC-I (p=0.523) downregulation.

To examine this relationship with proviral survival time at the intraparticipant level, we examined the 12 of the 16 participants in this study who had data for three or more OGV-derived *nef* clones. Assessing each participant individually by Spearman correlation, we only observed a significant relationship between CD4 downregulation and estimated proviral survival time of the corresponding *nef* variant (p<0.05), although this did not remain significant after p-value cutoff adjustment by Bonferroni correction (p<0.002) (**Table 2**). Linear regression analysis was also performed with discordant results (**Supplementary Table S2, Figures S3**, and **S4**). Overall, we found no convincing association between Nef function and the estimated proviral survival time of OGVs.

**Table 2.** Within-participant Spearman correlations between CD4 or MHC-I downregulation and estimated proviral survival time.

| 002 PID | Marker downregulated | n | Spearman's r (correlation co-efficient) | Spearman P-value (uncorrected) |
| --- | --- | --- | --- | --- |
| CAP188 | CD4 | 9 | 0.0667 | 0.8801 |
| CAP206 | CD4 | 9 | 0.5500 | 0.1328 |
| CAP217 | CD4 | 7 | -0.1786 | 0.7131 |
| CAP257 | CD4 | 7 | 0.5357 | 0.2357 |
| CAP268 | CD4 | 5 | 0.8000 | 0.1333 |
| CAP280 | CD4 | 5 | -0.1000 | 0.9500 |
| CAP287 | CD4 | 5 | -1.0000 | **0.0167\*** |
| CAP288 | CD4 | 5 | 0.6000 | 0.3500 |
| CAP302 | CD4 | 6 | -0.7714 | 0.1028 |
| CAP316 | CD4 | 6 | 0.4857 | 0.3556 |
| CAP336 | CD4 | 7 | 0.5714 | 0.2000 |
| CAP372 | CD4 | 6 | -0.9429 | **0.0167\*** |
| CAP188 | MHC-I | 9 | 0.1500 | 0.7081 |
| CAP206 | MHC-I | 9 | 0.3833 | 0.3125 |
| CAP217 | MHC-I | 7 | -0.2143 | 0.6615 |
| CAP257 | MHC-I | 7 | -0.7143 | 0.0881 |
| CAP268 | MHC-I | 5 | -0.2000 | 0.7833 |
| CAP280 | MHC-I | 5 | -0.7000 | 0.2333 |
| CAP287 | MHC-I | 5 | 0.9000 | 0.0833 |
| CAP288 | MHC-I | 5 | -0.1000 | 0.9500 |
| CAP302 | MHC-I | 6 | -0.7714 | 0.1028 |
| CAP316 | MHC-I | 6 | -0.8286 | 0.0583 |
| CAP336 | MHC-I | 7 | -0.1429 | 0.7825 |
| CAP372 | MHC-I | 6 | 0.4286 | 0.4194 |

## Discussion

Understanding the role of viral factors in the establishment and maintenance of the HIV reservoir is imperative to inform HIV cure strategies. We investigated the association between Nef function and reservoir proviral persistence in South African women living with HIV who initiated treatment in chronic infection. To our knowledge, this study is one of the first to investigate the function of *nef* variants obtained from HIV reservoir outgrowth viruses, and is the first to examine this in women living with HIV subtype C, the most prominent viral subtype worldwide [36]. In keeping with previous findings [37], we observed a relatively narrow functional range for CD4 downregulation by different *nef* variants within a participant despite *nef* sequence pairwise diversity of up to 5%, while there were greater differences in within-participant MHC-I downregulation.

We found a significant relationship between maximal MHC-I downregulation and the frequency of latently infected cells giving rise to viral outgrowth, which supports the earlier finding from North America that the function of plasma RNA-derived *nef* clones correlated positively with both HIV proviral DNA load and RC-VR size [31]. However, this association did not hold when adjusting for co-variates such as age and nadir CD4 count, the latter of which is a significant correlate of reservoir size [38] and timing of variant seeding [18] in our cohort. Furthermore, our study has several unique advantages. The first of these is making use of *nef* clones derived directly from the latent reservoir as opposed to plasma RNA-derived *nef* sequences. In addition, having timing estimates of reservoir entry for each outgrowth virus allowed us to estimate proviral ‘age’ at the time of reservoir sampling and explore a potential role for Nef in the HIV persistence. Here we observed little evidence of a survival benefit for proviral variants with stronger Nef MHC-I downregulation capacity. While our dataset included *nef* sequences estimated to enter the reservoir at various time-points pre-ART, we previously reported a distinct bias in the timing of entry to the year prior to ART initiation in individuals who initiated ART in chronic infection [16,18]. As a result, viruses seeded into the reservoir in acute/early stages of infection were under-represented, potentially precluding our ability to fully explore the role of Nef in proviral survival.

A second advantage is that our quantitative viral outgrowth assays were performed after the women had been on treatment for a median of five years compared to the 48-week post-treatment initiation sampling performed by Omondi *et al*. The benefit of our sampling is that our measurements were not impacted by the initial reservoir decay that occurs within the first two years after treatment initiation [39–41], resulting in a more representative sampling of the long-term RC-VR. Finally, a third advantage is that evaluating this question in the context of chronic ART initiation represents the majority of treatment initiations globally, representing a more real-world context.

Mechanisms of reservoir maintenance and persistence described thus far include homeostatic proliferation of infected reservoir cells [42], viral load blips, and low-level viraemia [43]. Definitive viral drivers of reservoir maintenance have not yet been completely elucidated. Nef proteins encoded by replication-competent viruses may contribute to reservoir persistence by facilitating the evasion of host CTLs through downregulation of cell surface MHC-class-I. In turn, sheltering cells harbouring proviruses containing intact *nef* genes. Studies have shown an enrichment of intact *nef* genes in proviral genomes from the reservoir [28,44], pointing to a role in viral persistence.

While we attempted to characterize as many Nef proteins as possible, using sequences obtained from a quantitative viral outgrowth assay has limitations. Outgrowth viruses represent only a portion of the intact latent reservoir [45] and preclude the analysis of *nef* variants from intact viruses that were not reactivated after one round of maximal stimulation *ex vivo*. Additionally, they do not account for *nef* variants that are potentially functional but located within defective proviruses, and while these Nef proteins are not directly associated with persistence of replication-competent proviruses, their ongoing expression may shape the proviral landscape over time and may contribute to ongoing immune activation [28,46].

This study focused on PLWH in a majority subtype C background, adding to the information on persistence in a non-B setting. This is pertinent as Nef function varies across HIV subtypes [46–48] and thus the resulting impact of Nef function on the latent reservoir may also differ across subtypes. While the scope of this study did not include comparing the function of Nef across subtypes, our study adds to the growing body of literature elucidating the role of Nef in modulating the RC-VR in different settings. Further studies with more diverse HIV subtype distributions should be pursued to examine this question.

Finally, ongoing maintenance of the viral reservoir is a multifaceted process that involves several viral, immunologic, and environmental factors, which makes it difficult to assess a specific effect size for one single attribute. However, given this complexity, our results identified a possible role for Nef-mediated MHC-I in this process, which agrees with previous work from unrelated cohorts. Taken together, these data support the further examination of the role of Nef in reservoir formation and maintenance, and as a possible target for therapeutic intervention toward the goal of an HIV cure.

## Materials and methods

### Study approval

This study was approved by the University of Cape Town Human Research Ethics Committee (718/2020), the University of KwaZulu Natal (BE178/150), the University of North Carolina at Chapel Hill (IRB 15-2717). All participants provided written informed consent prior to inclusion in the study.

### Study participants

The sixteen women included in this study were from the Centre of the AIDS Programme of Research in South Africa (CAPRISA) 002 acute infection cohort [49]. Quantitative viral outgrowth assays and measurement of T cell activation by flow cytometry have been described previously [18,50]. Briefly, resting CD4+ T cells were isolated from cryopreserved PBMCs obtained at a median of 5 years (IQR: 4.7-5.5; range: 4.1-7.3 years) after ART initiation. Bias-corrected maximum likelihood estimates for infectious units per million resting CD4^+^ T-cells (IUPM) were calculated in R using the SLDAssay package [51] based on the frequency of HIV-1 p24 capsid-positive wells on day 15 of the assay.

### nef sequencing, phylogenetic analysis, and selection of clones for assessment

*nef* sequences were derived from near full-length genome PacBio sequencing of viral variants from the QVOAs as previously described [16,18] and the resulting sequences have previously been deposited in GenBank (accession nos. MN097551 to MN097697 and OQ551935 to OQ552532). For each outgrowth virus variant, previously reported estimated reservoir entry timing [16,18] was used to calculate proviral survival time (weeks pre-ART seeded + time on ART). MLE amino acid trees were generated using PhyML version 3.3.20220420 [52] in DIVEIN with default settings [53]. DNA distances were calculated in MEGA 11 (version 11.0.13) [34] using a Maximum Composite Likelihood model (Poisson model with uniform substitution rates among sites and pairwise deletion of gaps) [54]. For examination in this study, *nef* genes were selected (i) that were phylogenetically distinct from one another, and (ii) such that a range of seeding times were included for each participant. Participant-specific *nef* genes were synthesized and cloned into the pNL4.3 ΔGag/Pol eGFP vector by GENEWIZ (Azenta Life Sciences, MA, USA).

### Cell culture

#### Pseudovirus generation and infections

Nef-typed pseudoviruses were produced by transfection of HEK-293T cells (ATCC CRL-3216) [55,56] as described previously [57]. Briefly, 1x10^6^ cells per well were seeded into a 6-well plate. Plasmid pNL4.3 ΔGag/Pol eGFP [57,58] containing the participant-specific *nef* gene, pCMV-DR8.2 (encoding Gag/Pol; Addgene catalogue number: 12263), and pMD2.G (encoding VSV-G; Addgene catalogue number: 12259) were co-transfected at a ratio of 0.4:1:1, respectively, as previously described [33]. 5 µg of plasmid mix was incubated with Lipofectamine 3000 transfection reagent (Thermo Scientific, USA) according to the manufacturer’s instructions. Thereafter, the transfection mix was added dropwise to each well of cells. The culture medium in each well was replaced after 24 hours. After a subsequent 48-hour incubation, supernatants were harvested, supplemented with 20% FBS, clarified by centrifugation at 500 x *g* for 5 minutes at room temperature, passed through a 0.45 µM cellulose acetate syringe filter, and aliquoted into cryotubes for storage at -80°C until needed.

For infection of Sup-T1 cells (ATCC CRL-1942) [59–64], pseudovirus aliquots were diluted to achieve 10 to 60 percent infection (as measured by the frequency of eGFP-positive Sup-T1 cells on day 2) and supplemented with 8 µg/mL hexadimethrine bromide (Sigma-Aldrich, MO, USA). 1x10^6^ Sup-T1 cells per well of a 24-well plate were pelleted (500 x *g* for 5 minutes at room temperature) and resuspended in the pseudovirus mix. Following 8 hours of incubation at 37°C in the presence of 5.5% CO_2_, the contents of each well were pelleted once again and resuspended in 1 mL complete RPMI, followed by incubation for a further 40 hours. All infections were performed in triplicate, and each experiment was repeated a minimum of three times. ΔNef negative and NL4-3 Nef positive control infections were included in triplicate in each experiment.

### Cell surface staining and flow cytometry

To measure downregulation of cell surface CD4 and MHC-I, Sup-T1 cells were collected at 48 hours after infection, pelleted (500 x *g* for 5 minutes) and transferred to wells of a V-bottom 96-well plate. Infected cells were washed twice with PBS, stained with LIVE/DEAD fixable Near-Infrared stain (Invitrogen) for 15 minutes at room temperature in the dark to identify dead cells. Sup-T1s were washed twice with FACS wash (PBS supplemented with 1% FCS; Capricorn Scientific) and subsequently stained with a cocktail of anti-human CD4-APC (BioLegend; clone OKT4) and anti-human HLA-A,B,C-BV605 (BioLegend; clone W6/32) for 20 minutes at room temperature in the dark. Finally, cells were washed three times with FACS wash, and fixed with Cell Fix (BD Biosciences). All flow cytometric data were acquired on a BD Fortessa and analysed using FlowJo v10.5.3 Software (BD Life Sciences). The gating strategy for downstream analysis is presented in **Figure S2**.

### Statistical analysis

Spearman rank correlation tests and area under the curve calculations for the VL (AUC VL) used in correlations and linear regression models for individual participants were performed in Prism version 8 (GraphPad). Linear regression and mixed effects models were performed in R (version 4.3.0). A P-value below 0.05 was considered statistically significant for reporting.

## Acknowledgements

The authors would like to thank the participants in the CAPRISA 002 acute infection cohort for their selfless contribution to HIV cure research. We’d also like to thank the CAPRISA study team and staff. Dr Cathrine Riou is also appreciated for providing flow cytometry guidance.

## Author email addresses

SDI

SS

BA

MCN

MJM

JDD

SBJ

MM

RS

NG

CW

TCQ

MRA

ADR

## Author contributions

ADR, MRA, SDI, MJM, and JDD conceived and designed experiments. In addition, JDD and MJM provided the reagent constructs, and the training required to carry out these experiments. SDI, BA, SS, SBJ, MM conducted experiments. SDI, MN performed statistical analyses. NG and CW assisted in oversight of the original CAPRISA study. TCQ, CW, and RS provided project guidance. SDI, MRA, and ADR aided in writing the manuscript. All authors read, edited, and approved the final manuscript.

## Data availability statement

The authors confirm that the data supporting the findings of this study are available within the manuscript and its Supporting Information files.

**Table S1. Nef subtype prediction for each OGV sequence using the Geno2Pheno Virus Detection and Subtyping tool.**

**Table S2. Within-participant linear regression analysis for the relationship between CD4 or MHC-I downregulation and estimated proviral survival time.**

---

**Figure S1.**
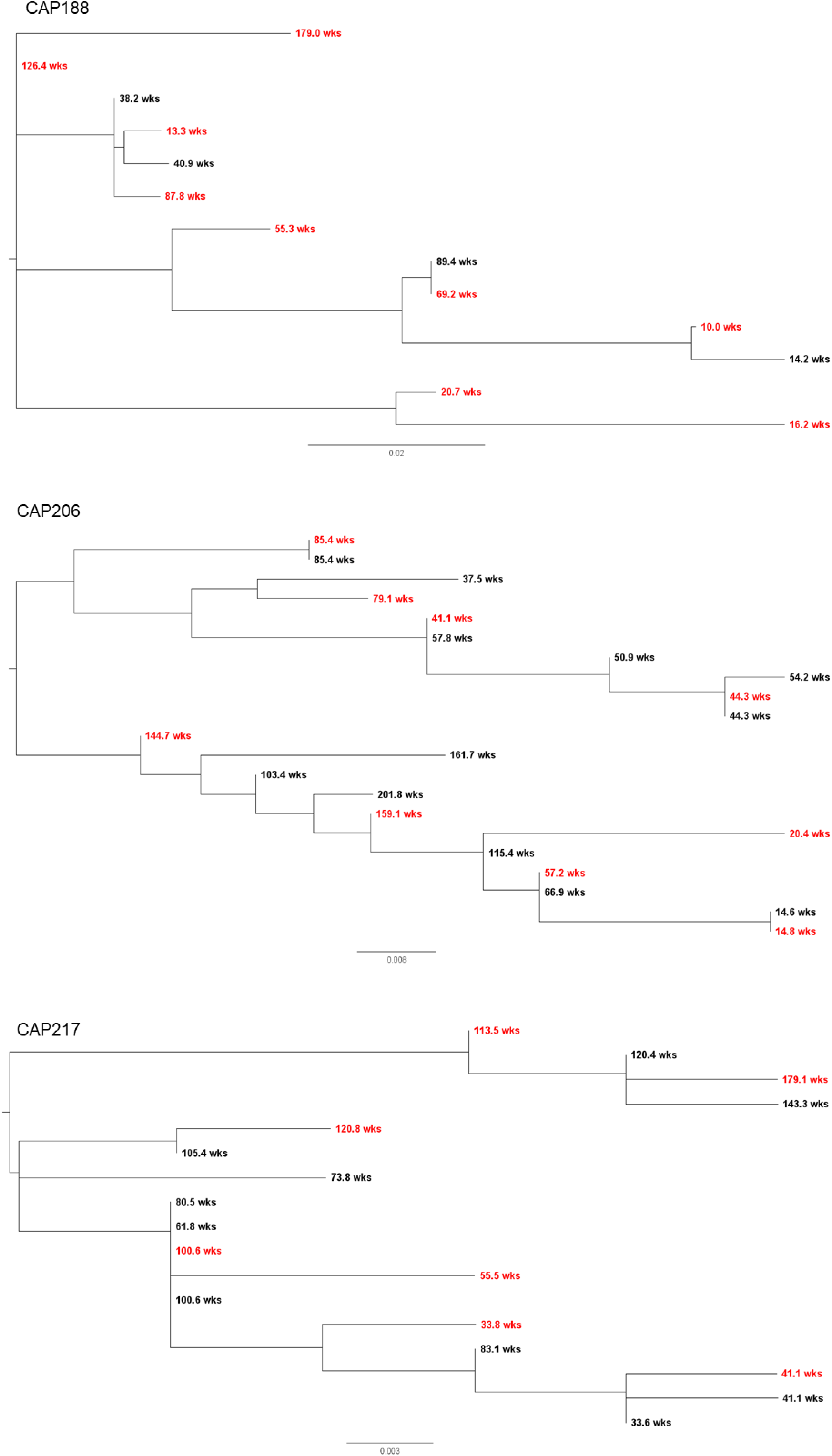
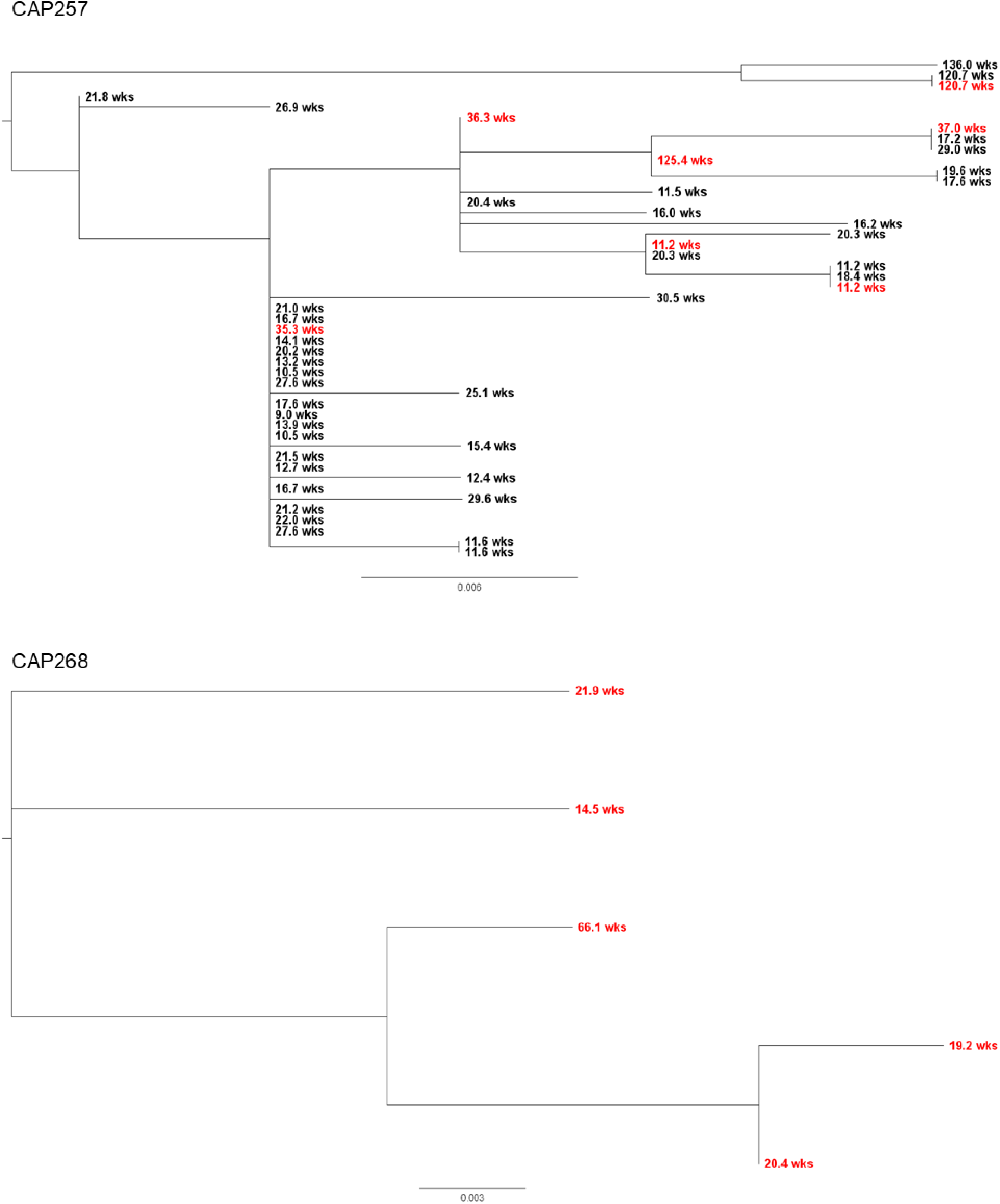
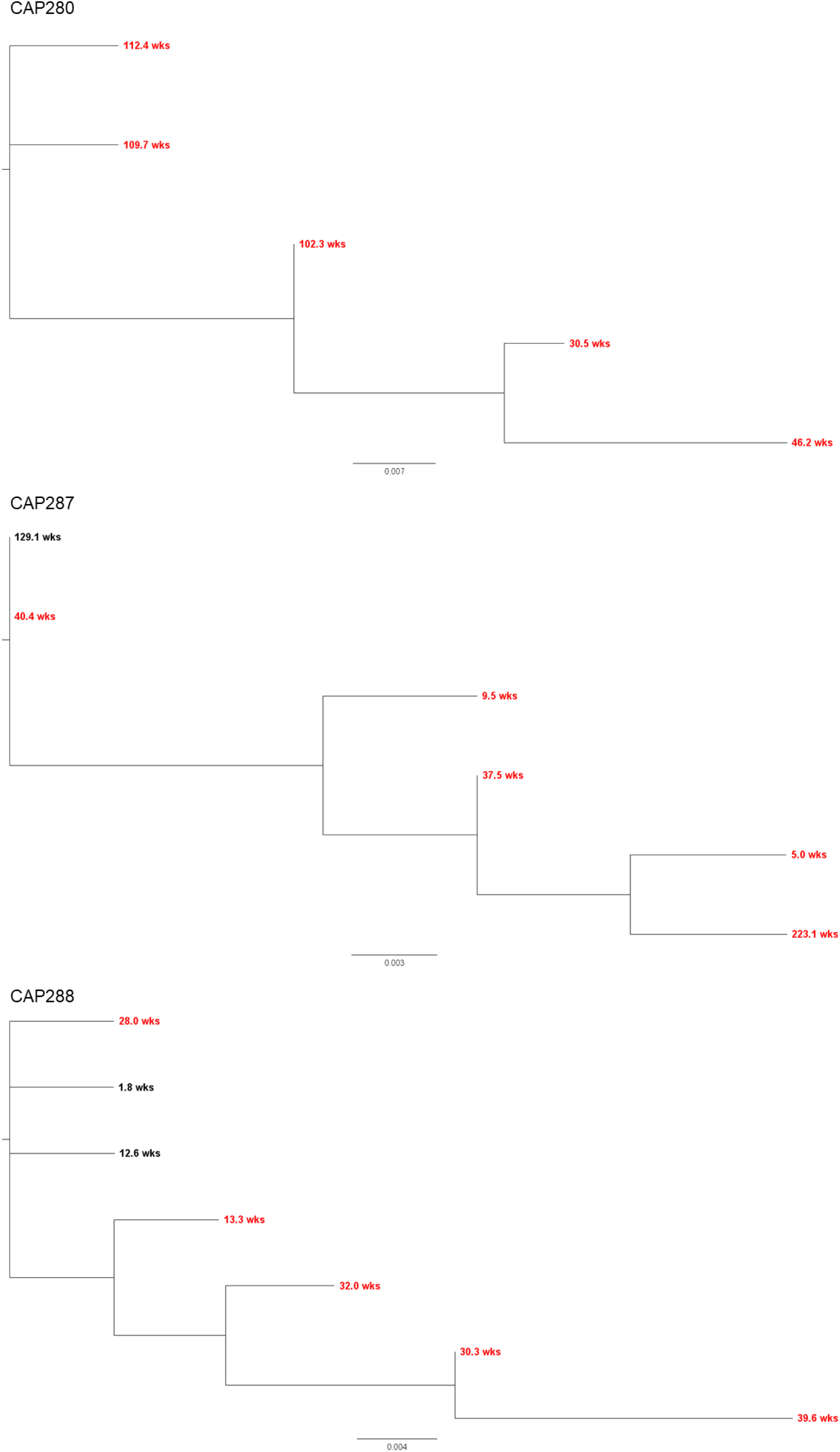
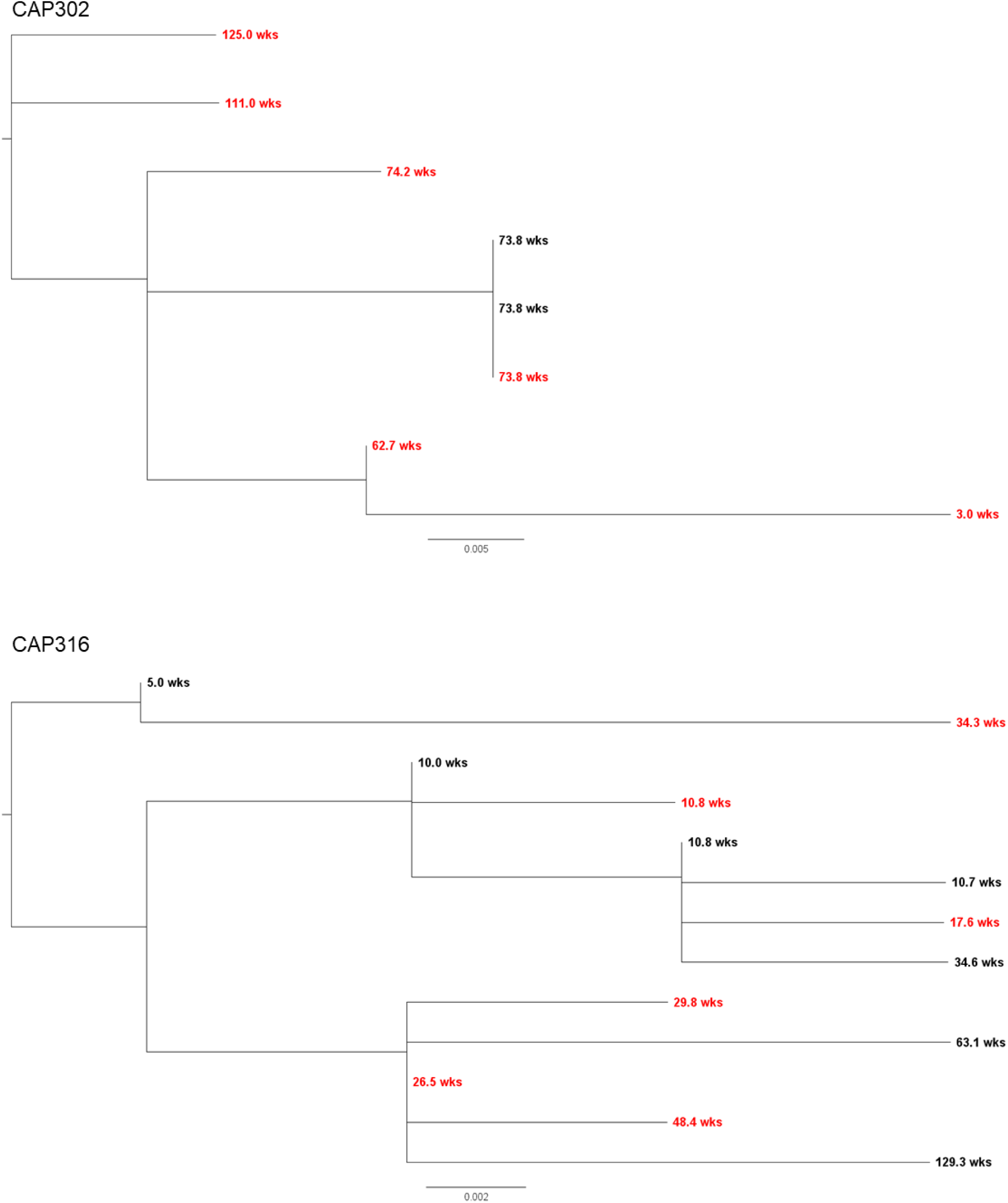
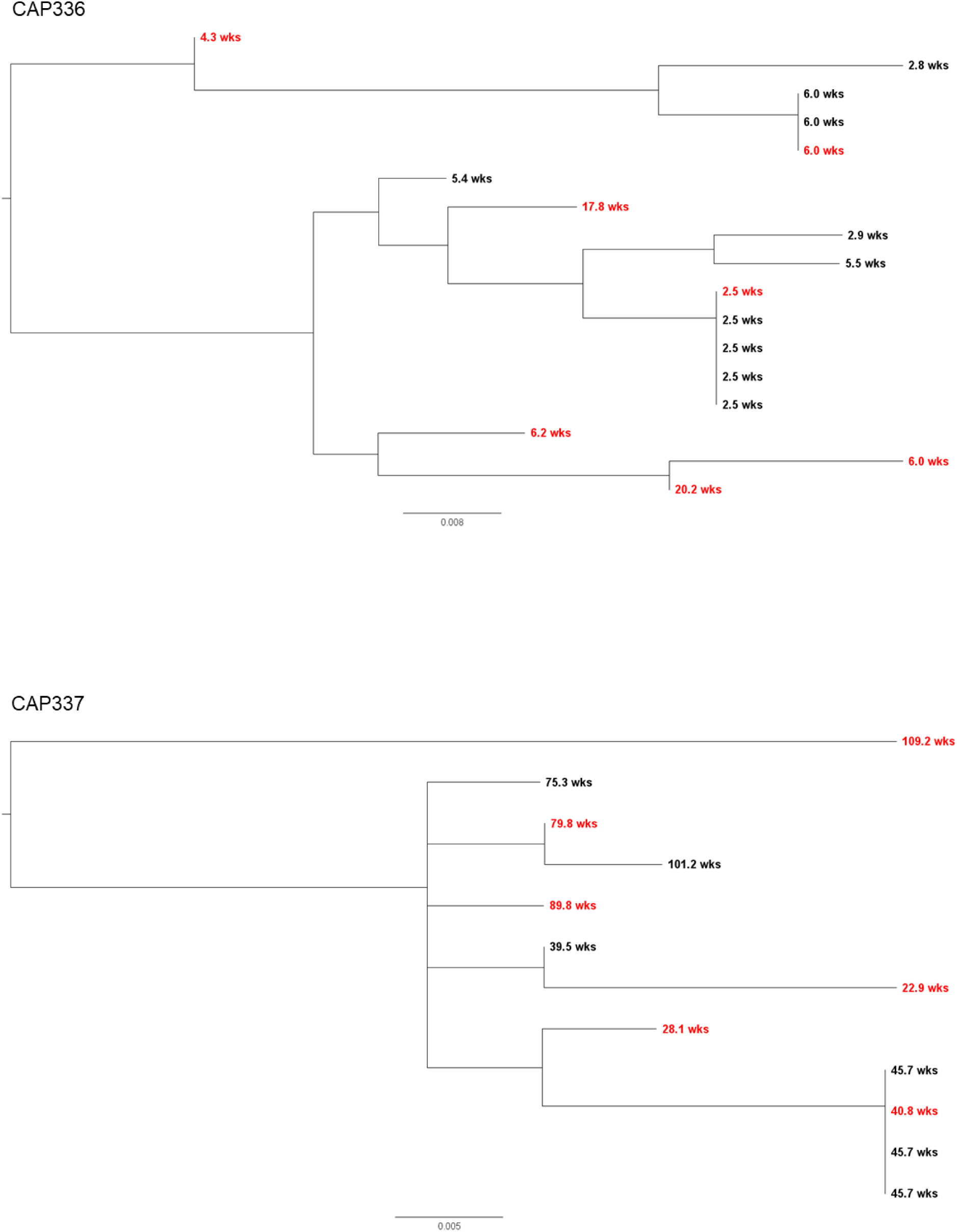
Individual participant Nef amino acid trees. OGV *nef* sequences were translated, aligned by Clustal W in MEGA11 [34] and trees were generated using PhyML [52] in DIVEIN [53]. Trees were mid-point rooted and sorted by decreasing node order. A scale bar is included below each tree. Labels indicate the time pre-ART that each OGV was estimated to enter the reservoir in weeks (wks). Red node labels indicate amino acid sequences that were selected for synthesis and subsequent testing for function. Black node labels represent OGV sequences that were not selected for further functional testing in this study.

**Figure S2.**
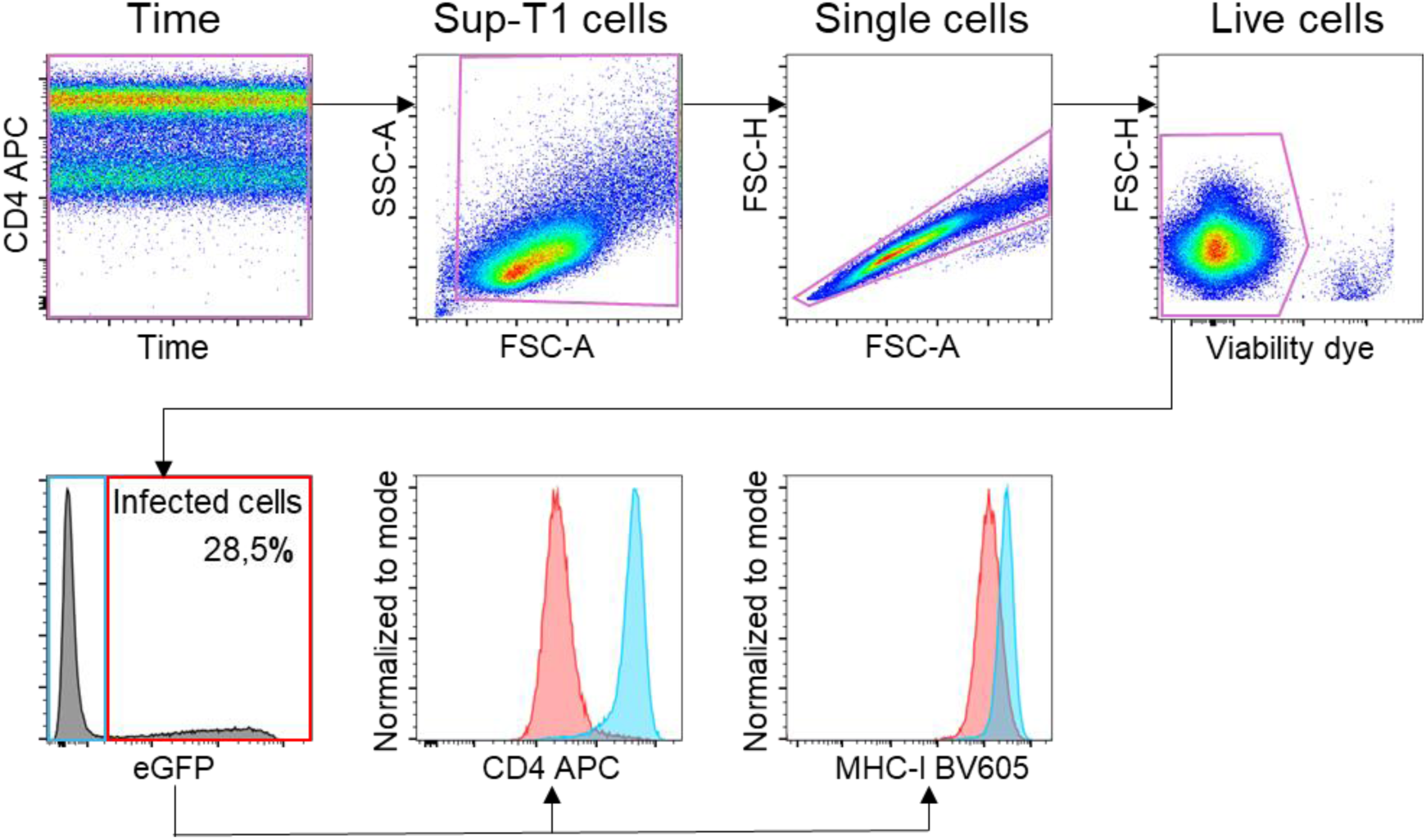
Gating strategy. Live infected Sup-T1 cells were infected with Nef-pseudotyped viruses. Sup-T1 cells were identified, and doublets were excluded before gating for live cells (gates indicated in purple) which were Near-Infrared^-^ (viability dye). Subsequently, infected cells were gated as eGFP^+^ events (red gate) while uninfected cells were identified as eGFP^-^ (blue gate). In this representative plot, 28.5% of live, single cells were infected. Gates were set based on single stained Sup-T1 cells as well as fluorescence minus one controls (FMOs), stained for all **fluorophores except that which is being gated on. Histogram overlays for either CD4 or MHC-I** expression show the fluorescence intensity shift between infected cells (red peaks) and uninfected cells (blue peaks).

**Figure S3.**
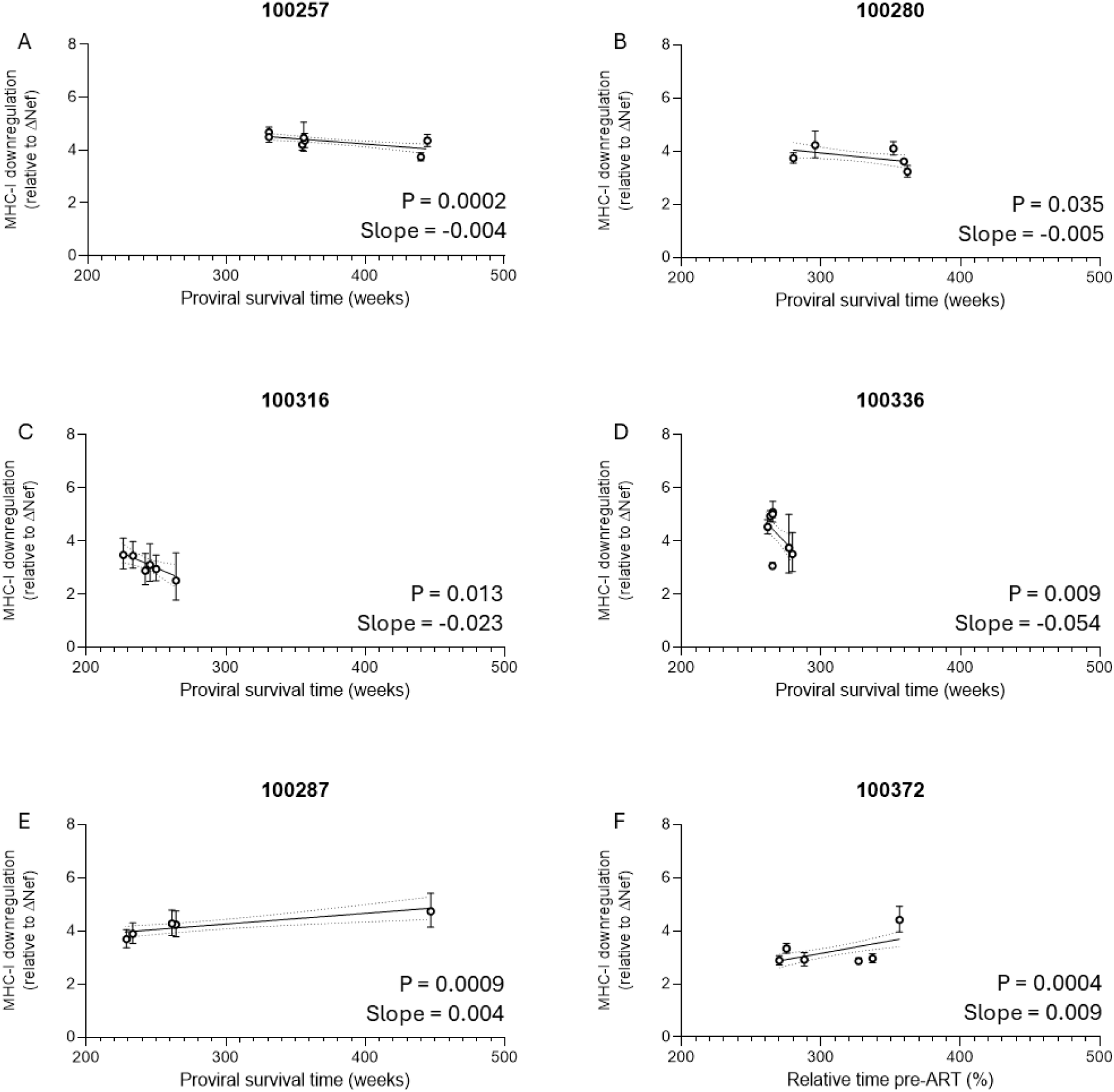
Participants with a significant linear relationship between MHC-I downregulation activity and proviral survival time. Each point on the graph represents geometric mean MHC-I downregulation for a unique *nef* clone and the error bars represent the 95% CI. Linear regression best fit lines and 95% CI were plotted for significant linear relationships.

**Figure S4.**
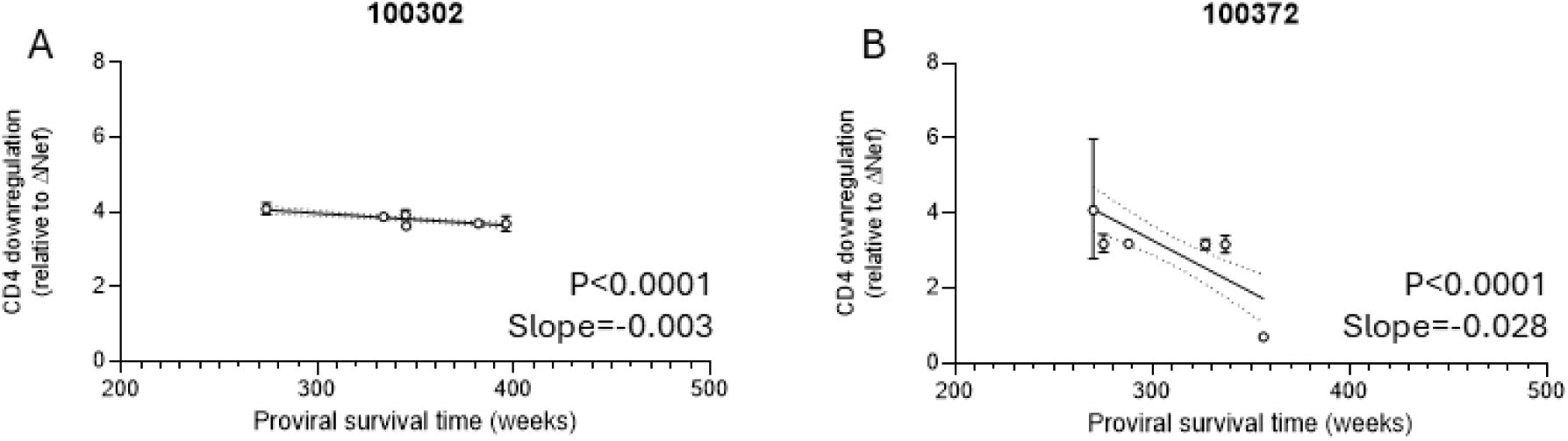
Individual participants with a significant linear relationship between CD4 downregulation activity and proviral survival time. Each point on the graph represents geometric mean CD4 downregulation for each *nef* clone and the error bars represent the 95% CI. Linear regression best fit lines and 95% CI were plotted for significant linear relationships.

## References

1. Finzi D. Identification of a Reservoir for HIV-1 in Patients on Highly Active Antiretroviral Therapy. Science. 1997;278: 1295–1300. doi:10.1126/science.278.5341.1295.

2. Finzi D, Blankson J, Siliciano JD, Margolick JB, Chadwick K, Pierson T, et al. Latent infection of CD4+ T cells provides a mechanism for lifelong persistence of HIV-1, even in patients on effective combination therapy. Nat Med. 1999;5: 512–517. doi:10.1038/8394.

3. Chun T-W, Stuyver L, Mizell SB, Ehler LA, Mican JAM, Baseler M, et al. Presence of an inducible HIV-1 latent reservoir during highly active antiretroviral therapy. Proc Natl Acad Sci. 1997;94: 13193–13197. doi:10.1073/pnas.94.24.13193.

4. Bruner KM, Murray AJ, Pollack RA, Soliman MG, Laskey SB, Capoferri AA, et al. Defective proviruses rapidly accumulate during acute HIV-1 infection. Nat Med. 2016;22: 1043–1049. doi:10.1038/nm.4156.

5. Jubault V, Burgard M, Le Corfec E, Costagliola D, Rouzioux C, Viard JP. High rebound of plasma and cellular HIV load after discontinuation of triple combination therapy. AIDS. 1998;12: 2358–9.

6. Neumann AU, Tubiana R, Calvez V, Robert C, Li TS, Agut H, et al. HIV-1 rebound during interruption of highly active antiretroviral therapy has no deleterious effect on reinitiated treatment. Comet Study Group. AIDS. 1999;13: 677–83. doi:10.1097/00002030-199904160-00008.

7. García F, Plana M, Vidal C, Cruceta A, O’Brien WA, Pantaleo G, et al. Dynamics of viral load rebound and immunological changes after stopping effective antiretroviral therapy. AIDS. 1999;13: F79–86. doi:10.1097/00002030-199907300-00002.

8. Davey RT, Bhat N, Yoder C, Chun T-W, Metcalf JA, Dewar R, et al. HIV-1 and T cell dynamics after interruption of highly active antiretroviral therapy (HAART) in patients with a history of sustained viral suppression. Proc Natl Acad Sci. 1999;96: 15109–15114. doi:10.1073/pnas.96.26.15109.

9. Ioannidis JP, Havlir D V, Tebas P, Hirsch MS, Collier AC, Richman DD. Dynamics of HIV-1 viral load rebound among patients with previous suppression of viral replication. AIDS. 2000;14: 1481–8. doi:10.1097/00002030-200007280-00003.

10. Whitney JB, Hill AL, Sanisetty S, Penaloza-MacMaster P, Liu J, Shetty M, et al. Rapid seeding of the viral reservoir prior to SIV viraemia in rhesus monkeys. Nature. 2014;512: 74–77. doi:10.1038/nature13594.

11. Buzon MJ, Martin-Gayo E, Pereyra F, Ouyang Z, Sun H, Li JZ, et al. Long-Term Antiretroviral Treatment Initiated at Primary HIV-1 Infection Affects the Size, Composition, and Decay Kinetics of the Reservoir of HIV-1-Infected CD4 T Cells. J Virol. 2014;88: 10056–10065. doi:10.1128/JVI.01046-14.

12. Ananworanich J, Schuetz A, Vandergeeten C, Sereti I, de Souza M, Rerknimitr R, et al. Impact of Multi-Targeted Antiretroviral Treatment on Gut T Cell Depletion and HIV Reservoir Seeding during Acute HIV Infection. PLoS One. 2012;7: e33948. doi:10.1371/journal.pone.0033948.

13. Hocqueloux L, Avettand-Fènoël V, Jacquot S, Prazuck T, Legac E, Mélard A, et al. Long-term antiretroviral therapy initiated during primary HIV-1 infection is key to achieving both low HIV reservoirs and normal T cell counts. J Antimicrob Chemother. 2013;68: 1169–1178. doi:10.1093/jac/dks533.

14. Brodin J, Zanini F, Thebo L, Lanz C, Bratt G, Neher RA, et al. Establishment and stability of the latent HIV-1 DNA reservoir. Elife. 2016;5. doi:10.7554/eLife.18889.

15. Jones BR, Kinloch NN, Horacsek J, Ganase B, Harris M, Harrigan PR, et al. Phylogenetic approach to recover integration dates of latent HIV sequences within-host. Proc Natl Acad Sci. 2018;115: E8958–E8967. doi:10.1073/pnas.1802028115.

16. Abrahams M-R, Joseph SB, Garrett N, Tyers L, Moeser M, Archin N, et al. The replication-competent HIV-1 latent reservoir is primarily established near the time of therapy initiation. Sci Transl Med. 2019;11: eaaw5589. doi:10.1126/scitranslmed.aaw5589.

17. Pankau MD, Reeves DB, Harkins E, Ronen K, Jaoko W, Mandaliya K, et al. Dynamics of HIV DNA reservoir seeding in a cohort of superinfected Kenyan women. PLoS Pathog. 2020;16: e1008286. doi:10.1371/journal.ppat.1008286.

18. Joseph SB, Abrahams M-RR, Moeser M, Tyers L, Archin NM, Council OD, et al. The timing of HIV-1 infection of cells that persist on therapy is not strongly influenced by replication competency or cellular tropism of the provirus. PLoS Pathog. 2024;20: e1011974. doi:10.1371/journal.ppat.1011974.

19. Buffalo CZ, Iwamoto Y, Hurley JH, Ren X. How HIV Nef Proteins Hijack Membrane Traffic To Promote Infection. J Virol. 2019;93. doi:10.1128/JVI.01322-19.

20. Ross TM, Oran AE, Cullen BR. Inhibition of HIV-1 progeny virion release by cell-surface CD4 is relieved by expression of the viral Nef protein. Curr Biol. 1999;9: 613–621. doi:10.1016/S0960-9822(99)80283-8.

21. Argañaraz ER, Schindler M, Kirchhoff F, Cortes MJ, Lama J. Enhanced CD4 Down-modulation by Late Stage HIV-1 nef Alleles Is Associated with Increased Env Incorporation and Viral Replication. J Biol Chem. 2003;278: 33912–33919. doi:10.1074/jbc.M303679200.

22. Prévost J, Richard J, Medjahed H, Alexander A, Jones J, Kappes JC, et al. Incomplete Downregulation of CD4 Expression Affects HIV-1 Env Conformation and Antibody-Dependent Cellular Cytotoxicity Responses. J Virol. 2018;92: 1–16. doi:10.1128/JVI.00484-18.

23. Schwartz O, Maréchal V, Gall S Le, Lemonnier F, Heard J-M. Endocytosis of major histocompatibility complex class I molecules is induced by the HIV–1 Nef protein. Nat Med. 1996;2: 338–342. doi:10.1038/nm0396-338.

24. Collins KL, Chen BK, Kalams SA, Walker BD, Baltimore D. HIV-1 Nef protein protects infected primary cells against killing by cytotoxic T lymphocytes. Nature. 1998;391: 397–401. doi:10.1038/34929.

25. Ferdin J, Goričar K, Dolžan V, Plemenitaš A, Martin JN, Peterlin BM, et al. Viral protein Nef is detected in plasma of half of HIV-infected adults with undetectable plasma HIV RNA. PLoS One. 2018;13: 1–10. doi:10.1371/journal.pone.0191613.

26. Wang T, Green LA, Gupta SK, Amet T, Byrd DJ, Yu Q, et al. Intracellular Nef Detected in Peripheral Blood Mononuclear Cells from HIV Patients. AIDS Res Hum Retroviruses. 2015;31: 217–220. doi:10.1089/aid.2013.0250.

27. Stevenson EM, Ward AR, Truong R, Thomas AS, Huang S-H, Dilling TR, et al. HIV-specific T cell responses reflect substantive in vivo interactions with antigen despite long-term therapy. JCI Insight. 2021;6. doi:10.1172/jci.insight.142640.

28. Duette G, Hiener B, Morgan H, Mazur FG, Mathivanan V, Horsburgh BA, et al. The HIV-1 proviral landscape reveals that Nef contributes to HIV-1 persistence in effector memory CD4+ T cells. J Clin Invest. 2022;132. doi:10.1172/JCI154422.

29. Pollack RA, Jones RB, Pertea M, Bruner KM, Martin AR, Thomas AS, et al. Defective HIV-1 Proviruses Are Expressed and Can Be Recognized by Cytotoxic T Lymphocytes, which Shape the Proviral Landscape. Cell Host Microbe. 2017;21: 494–506.e4. doi:10.1016/j.chom.2017.03.008.

30. Imamichi H, Smith M, Adelsberger JW, Izumi T, Scrimieri F, Sherman BT, et al. Defective HIV-1 proviruses produce viral proteins. Proc Natl Acad Sci U S A. 2020;117: 3704–3710. doi:10.1073/pnas.1917876117.

31. Omondi FH, Chandrarathna S, Mujib S, Brumme CJ, Jin SW, Sudderuddin H, et al. HIV Subtype and Nef-Mediated Immune Evasion Function Correlate with Viral Reservoir Size in Early-Treated Individuals. J Virol. 2019;93. doi:10.1128/JVI.01832-18.

32. Pirkl M, Büch J, Friedrich G, Böhm M, Turner D, Degen O, et al. Geno2pheno: recombination detection for HIV-1 and HEV subtypes. NAR Mol Med. 2024;1: 1–9. doi:10.1093/narmme/ugae003.

33. Pawlak EN, Dirk BS, Jacob RA, Johnson AL, Dikeakos JD. The HIV-1 accessory proteins Nef and Vpu downregulate total and cell surface CD28 in CD4+ T cells. Retrovirology. 2018;15: 6. doi:10.1186/s12977-018-0388-3.

34. Tamura K, Stecher G, Kumar S. MEGA11: Molecular Evolutionary Genetics Analysis Version 11. Mol Biol Evol. 2021;38: 3022–3027. doi:10.1093/molbev/msab120.

35. Saitou N, Nei M. The neighbor-joining method: a new method for reconstructing phylogenetic trees. Mol Biol Evol. 1987;4: 406–425. doi:10.1093/oxfordjournals.molbev.a040454.

36. Bbosa N, Kaleebu P, Ssemwanga D. HIV subtype diversity worldwide. Curr Opin HIV AIDS. 2019;14: 153–160. doi:10.1097/COH.0000000000000534.

37. Mwimanzi P, Markle TJ, Ogata Y, Martin E, Tokunaga M, Mahiti M, et al. Dynamic range of Nef functions in chronic HIV-1 infection. Virology. 2013;439: 74–80. doi:10.1016/j.virol.2013.02.005.

38. Ismail SD, Riou C, Joseph SB, Archin NM, Margolis DM, Perelson AS, et al. Immunological Correlates of the HIV-1 Replication-Competent Reservoir Size. Clin Infect Dis. 2021;73: 1528–1531. doi:10.1093/cid/ciab587.

39. Chun T-W, Justement JS, Moir S, Hallahan CW, Maenza J, Mullins JI, et al. Decay of the HIV reservoir in patients receiving antiretroviral therapy for extended periods: implications for eradication of virus. J Infect Dis. 2007;195: 1762–4. doi:10.1086/518250.

40. Archin NM, Vaidya NK, Kuruc JAD, Liberty AL, Wiegand A, Kearney MF, et al. Immediate antiviral therapy appears to restrict resting CD4 + cell HIV-1 infection without accelerating the decay of latent infection. Proc Natl Acad Sci U S A. 2012;109: 9523–9528. doi:10.1073/pnas.1120248109.

41. Peluso MJ, Bacchetti P, Ritter KD, Beg S, Lai J, Martin JN, et al. Differential decay of intact and defective proviral DNA in HIV-1–infected individuals on suppressive antiretroviral therapy. JCI Insight. 2020;5: 1–13. doi:10.1172/jci.insight.132997.

42. Chomont N, El-Far M, Ancuta P, Trautmann L, Procopio FA, Yassine-Diab B, et al. HIV reservoir size and persistence are driven by T cell survival and homeostatic proliferation. Nat Med. 2009;15: 893–900. doi:10.1038/nm.1972.

43. Bachmann N, von Siebenthal C, Vongrad V, Turk T, Neumann K, Beerenwinkel N, et al. Determinants of HIV-1 reservoir size and long-term dynamics during suppressive ART. Nat Commun. 2019;10: 3193. doi:10.1038/s41467-019-10884-9.

44. Dufour C, Richard C, Pardons M, Massanella M, Ackaoui A, Murrell B, et al. Phenotypic characterization of single CD4+ T cells harboring genetically intact and inducible HIV genomes. Nat Commun. 2023;14. doi:10.1038/s41467-023-36772-x.

45. Ho YC, Shan L, Hosmane NN, Wang J, Laskey SB, Rosenbloom DIS, et al. Replication-competent noninduced proviruses in the latent reservoir increase barrier to HIV-1 cure. Cell. 2013;155: 540–551. doi:10.1016/j.cell.2013.09.020.

46. Lamers SL, Fogel GB, Liu ES, Nolan DJ, Rose R, McGrath MS. HIV-1 subtypes maintain distinctive physicochemical signatures in Nef domains associated with immunoregulation. Infect Genet Evol. 2023;115. doi:10.1016/j.meegid.2023.105514.

47. Jin SW, Mwimanzi FM, Mann JK, Bwana MB, Lee GQ, Brumme CJ, et al. Variation in HIV-1 nef function within and among viral subtypes reveals genetically separable antagonism of serinc3 and serinc5. PLoS Pathog. 2020;16. doi:10.1371/journal.ppat.1008813.

48. Mann JK, Byakwaga H, Kuang XT, Le AQ, Brumme CJ, Mwimanzi P, et al. Ability of HIV-1 Nef to downregulate CD4 and HLA class I differs among viral subtypes. Retrovirology. 2013;10: 100. doi:10.1186/1742-4690-10-100.

49. van Loggerenberg F, Mlisana K, Williamson C, Auld SC, Morris L, Gray CM, et al. Establishing a cohort at high risk of HIV infection in South Africa: challenges and experiences of the CAPRISA 002 acute infection study. PLoS One. 2008;3: e1954. doi:10.1371/journal.pone.0001954.

50. Abrahams M-R, Anderson JA, Giorgi EE, Seoighe C, Mlisana K, Ping L-H, et al. Quantitating the Multiplicity of Infection with Human Immunodeficiency Virus Type 1 Subtype C Reveals a Non-Poisson Distribution of Transmitted Variants. J Virol. 2009;83: 3556–3567. doi:10.1128/JVI.02132-08.

51. Trumble IM, Allmon AG, Archin NM, Rigdon J, Francis O, Baldoni PL, et al. SLDAssay: A software package and web tool for analyzing limiting dilution assays. J Immunol Methods. 2017;450: 10–16. doi:10.1016/j.jim.2017.07.004.

52. Guindon S, Dufayard JF, Lefort V, Anisimova M, Hordijk W, Gascuel O. New algorithms and methods to estimate maximum-likelihood phylogenies: Assessing the performance of PhyML 3.0. Syst Biol. 2010;59: 307–321. doi:10.1093/sysbio/syq010.

53. Deng W, Maust BS, Nickle DC, Learn GH, Liu Y, Heath L, et al. DIVEIN: a web server to analyze phylogenies, sequence divergence, diversity, and informative sites. Biotechniques. 2010;48: 405–408. doi:10.2144/000113370.

54. Tamura K, Nei M, Kumar S. Prospects for inferring very large phylogenies by using the neighbor-joining method. Proc Natl Acad Sci. 2004;101: 11030–11035. doi:10.1073/pnas.0404206101.

55. DuBridge RB, Tang P, Hsia HC, Leong PM, Miller JH, Calos MP. Analysis of mutation in human cells by using an Epstein-Barr virus shuttle system. Mol Cell Biol. 1987;7: 379–387. doi:10.1128/MCB.7.1.379.

56. Pear WS, Nolan GP, Scott ML, Baltimore D. Production of high-titer helper-free retroviruses by transient transfection. Proc Natl Acad Sci U S A. 1993;90: 8392–6.

57. Mumby MJ, Johnson AL, Trothen SM, Edgar CR, Gibson R, Stathopulos PB, et al. An Amino Acid Polymorphism within the HIV-1 Nef Dileucine Motif Functionally Uncouples Cell Surface CD4 and SERINC5 Downregulation. J Virol. 2021;95. doi:10.1128/JVI.00588-21.

58. Husain M, Gusella GL, Klotman ME, Gelman IH, Ross MD, Schwartz EJ, et al. HIV-1 Nef Induces Proliferation and Anchorage-Independent Growth in Podocytes. J Am Soc Nephrol. 2002;13: 1806–1815. doi:10.1097/01.ASN.0000019642.55998.69.

59. Smith SD, Shatsky M, Cohen PS, Warnke R, Link MP, Glader BE. Monoclonal antibody and enzymatic profiles of human malignant T-lymphoid cells and derived cell lines. Cancer Res. 1984;44: 5657–60.

60. Hecht F, Morgan R, Hecht BK, Smith SD. Common Region on Chromosome 14 in T-Cell Leukemia and Lymphoma. Science. 1984;226: 1445–1447. doi:10.1126/science.6438800.

61. Baer R, Chen K-C, Smith SD, Rabbitts TH. Fusion of an immunoglobulin variable gene and a T cell receptor constant gene in the chromosome 14 inversion associated with T cell tumors. Cell. 1985;43: 705–713. doi:10.1016/0092-8674(85)90243-0.

62. Lifson JD, Reyes GR, McGrath MS, Stein BS, Engleman EG. AIDS Retrovirus Induced Cytopathology: Giant Cell Formation and Involvement of CD4 antigen. Science. 1986;232: 1123–1127. doi:10.1126/science.3010463.

63. Reynolds TC, Smith SD, Sklar J. Analysis of DNA surrounding the breakpoints of chromosomal translocations involving the β T cell receptor gene in human lymphoblastic neoplasms. Cell. 1987;50: 107–117. doi:10.1016/0092-8674(87)90667-2.

64. Smith S, Morgan R, Gemmell R, Amylon M, Link M, Linker C, et al. Clinical and biologic characterization of T-cell neoplasias with rearrangements of chromosome 7 band q34. Blood. 1988;71: 395–402. doi:10.1182/blood.V71.2.395.395.

